# Cortical layer-specific and cell type-specific dendritic expression of Arc (Arg 3.1) in an *in vitro* model of slow-wave sleep

**DOI:** 10.1101/2024.07.17.603872

**Authors:** Iain J. Hartnell, Anna Simon, Stephen P. Hall, Fiona Ducotterd, Miles A. Whittington, Sangeeta Chawla

**Author notes:** Corresponding author: Dr Sangeeta Chawla, Department of Biology and York Biomedical Research Institute, University of York, Wentworth Way, Heslington, York YO10 5DD, UK. Deceased 21 October 2021 (Miles A. Whittington).

## Abstract

An abundance of evidence shows sleep homeostatically regulates synaptic plasticity and memory consolidation but the underlying molecular regulators of this important function of sleep have been difficult to directly elucidate. Arc is an immediate early gene that has a fundamental role in several aspects of synaptic plasticity including regulation and maintenance. Using a physiological brain slice model of slow-wave sleep, here we show that Arc protein has a characteristic spatial and cell-type specific distribution during persistent sleep-related delta oscillations in neocortical slices, which may correlate with aspects of sleep-dependent regulation of synaptic plasticity in cortical regions. In delta oscillating slices, Arc is highly expressed in layer 2/3 dendrites of the primary and secondary association cortex and this dendritic localisation is specific to intrinsically bursting cells (IB) whose cell bodies are in layer 5. Moreover, Arc immunopositive dendrites are clustered together arranged in a quasi hexagonal arrangement with a spacing of ∼ 50 µm between clusters. The cytoarchitectural distribution of Arc across the association cortex has implications for the mechanisms of sleep and for the synaptic homeostasis hypothesis regarding the function of sleep.

## Introduction

Sleep is required throughout organismal life for many vital bodily functions such as development, immune responses, and cognition. Sleep is particularly important for memory consolidation [1]. As memory acquisition during the wake state depends on synaptic potentiation in many circuits, one proposed function of sleep is to rescale synapses to prevent both runaway synaptic potentiation and the saturation of circuits in their ability to undergo synaptic potentiation. This synaptic homeostasis hypothesis (SHY) regarding the function of sleep was proposed by Tononi and Cirelli [2] and posits that synaptic downscaling occurs in particular during slow-wave sleep (SWS) or NREM (non-rapid eye movement) phase of sleep.

There is considerable electrophysiological, biochemical and structural evidence that synaptic plasticity is downscaled during sleep in rodents. In the rodent cortex, evoked excitatory field potentials are larger in the wake state and decrease during sleep [3]. Sleep also decreases the expression of surface GluA1-containing AMPA receptors ([3]; [4]; [5]) and reduces synapse size [6]. However, whilst there is net downscaling across the cortex, some heterogeneity persists and not all synapses show downscaling during sleep. For example, structural scaling was not evident in large synapses lacking endosomes [6] and although cortical expression levels of the plasticity-associated genes Arc and BDNF were high during waking and low during sleep, there was considerable variation across cortical layers [7]. This heterogeneity could result from synaptic scaling that is proportional to the relative synaptic weights and would also support bidirectional scaling during sleep [8].

Slow wave sleep (SWS) is important for synaptic scaling and reduced SWS is reported in neurological disorders such as Alzheimer’s disease [9], and so it is important to understand the underlying molecular mechanisms linking SWS to synaptic plasticity. SWS is characterised by slow wave activity (SWA), synchronous low frequency oscillations in the 0.5-4 Hz range (delta oscillations) evident in cortical electroencephalogram (EEG) recordings [10]. High frequency EEG oscillations in the 30-80 Hz range (gamma oscillations), on the other hand, represent cortical dynamics associated with sensory perception during wakefulness aiding attentional processing [11] and working memory [12]. SHY proposes that synaptic downscaling is mechanistically linked to SWA (delta oscillations). However, the functional, molecular, and structural synaptic downscaling in the aforementioned studies involved measurements after periods of sleep ranging from 3 to 7 hours. These periods of sleep include REM sleep. Additionally, NREM sleep is characterised by other electrophysiological features including sleep spindles that are observed in thalamocortical circuits as 7–15 Hz oscillations [13]. Thus, it is unclear which electrophysiological features are important in the process of synaptic downscaling and how the changes are related to cortical network dynamics.

To better understand the relationship between sleep/wake-related cortical network activity and synaptic plasticity we compared Arc expression in rodent cortical slices exhibiting pharmacologically generated sleep (delta) and wake (gamma) rhythms. Arc protein levels have previously been shown to change between sleep/wake states [7]. Arc is an effector immediate early gene whose expression is tightly regulated by synaptic activity at the level of transcription [14] and local translation of synaptically localised Arc mRNA at activated synapses ([15]; [16]). Arc’s molecular functions have the potential to regulate synaptic downscaling: Arc mediates AMPA receptor endocytosis [17,18] and regulates neuronal structural plasticity through regulation of spine morphology [19]. A more recently identified function of Arc is its ability to package its own mRNA into viral-like capsids to mediate intercellular mRNA transfer through extracellular vesicles [20]. Since Arc expression is induced selectively in activated synapses, its presence could have functional consequences and some or all of Arc’s diverse molecular actions in AMPA receptor trafficking, spine remodelling and intercellular transfer of its own mRNA could be involved in sleep-related synaptic scaling.

Here we demonstrate a complex spatial profile of Arc protein expression across the cortical layers in an *in vitro* model of slow wave sleep network activity and show that dendritic Arc expression is high in layer 5 intrinsically bursting cells.

## Methods

### Electrophysiology

Coronal sections, 450 μm thick, containing primary sensory and adjacent secondary, association neocortex were prepared from brains of adult male Wistar rats (150-200g) following cardiac perfusion with ice-cold buffered sucrose aCSF (252mM sucrose, 3mM KCl, 1.25mM NaH_2_PO_4_, 24mM NaHCO_3_, 2mM MgSO_4_, 2mM CaCl_2_, 10mM glucose). Slices were maintained at 33°C at the interface chamber between ACSF (126mM NaCl, 3mM KCl, 1.25mM NaH_2_PO_4_, 24mM NaHCO_3_, 1mM MgSO_4_, 1.2mM CaCl_2_, 10mM glucose) and warm, wetted 95% O2/5% CO2. In some experiments tangential sections were taken from the neocortex at the level of layer 2/3. All animals were housed with a constant supply of food and water, and a 12-hour light-dark cycle. Slices were prepared 3-4 hours into the light cycle.

Local field potentials (LFP) were recorded from coronal sections using glass microelectrodes (resistance >1 MΩ) filled with aCSF. To generate slow wave sleep-related delta-frequency oscillations in the primary and secondary somatosensory cortices a combination of the acetylcholine agonist, Carbachol (4 µM) and dopaminergic antagonist, SCH23990 (10 µM) were simultaneously added to the circulating aCSF. Cortical LFP recordings were made from layer 5 of the primary and secondary somatosensory cortex in coronal slices as described previously [21]. To generate wake-related gamma-frequency oscillations in the primary auditory and secondary somatosensory cortex kainic acid was used (400nM) on horizontal slices. LFP recordings were made from layer 3 of the primary auditory and secondary somatosensory cortex. Slices were allowed to express either delta or gamma rhythms for 2h before preparation for immunocytochemistry.

All LFP recordings were analysed for oscillatory activity by Fast Fourier transformations to generate power spectra. From the power spectra, measures of the frequency and modal peak power were taken. Power was integrated from the lower to the upper boundaries of the frequency bands of interest to give area power measurements. Limits of the frequency band were set for calculations of power: for gamma oscillations, these were 30-50Hz, and for delta, these were 0.5-4 Hz.

Intracellular recordings were made with sharp electrodes with a resistance of 70-200 MΩ filled with a 2M potassium acetate and 2% biocytin hydrochloride. To visually highlight single neurons that had been previously characterised by intracellular recording, the cell was maintained in a slightly hyperpolarized state whilst a small amount of tuning current allowed the biocytin (in the electrode) to fill the cell and its projections. Each cell was filled for 30 – 60 minutes and when complete the slice was removed and fixed in 4% paraformaldehyde (PFA).

### Immunocytochemistry

Slices from electrophysiological studies were fixed in 4% PFA overnight followed by cryoprotection with sucrose solutions (30% Sucrose in PBS respectively). The resultant slices were then immersed in optimum cutting temperature compound (OCT) frozen in Isopentane (2-Methylbutane) in dry ice and re-sectioned to 30μm. Selected slices were then washed in PBS, and PBS-TX (with 10% Methanol for nuclear stains) for 10 mins and blocked with 3% normal goat serum (NGS) for 2 hours. Primary antibodies were then applied: Millipore ABN90 Anti NeuN (1:500), Millipore MAB5406 Anti Gad67 (mouse, 1:500), Synaptic Systems Gmbh 156003 Anti Arc (rabbit, 1:500). Secondary antibodies used were: Alexa Fluor Goat-α-Rabbit 488 (1:500), Alexa Fluor Goat-α-Mouse 546 (1:500) and : Alexa Fluor Goat-α-Guinea Pig 633 (1:500). Alexa Fluor 568 Streptavidin conjugate (1:500) was used to detect biocytin. Slices were mounted on slides and imaged using AxioScan Z1.

Higher resolution images were taken using a Zeiss LSM 710. Confocal images were taken as z-stacks with the maximum intensity projection calculated for each image presented.

### Image analysis

Intensity-matched images were separated into the 3 triple-stained channels and converted into 16 bit grayscale .tif files for further analysis in Matlab. Primary and secondary cortical subregions were identified according to Paxinos & Watson (1998) and templates created according to the thresholded intensity of NeuN or GAD67 signal. In each case 1 standard deviation from the mean intensity was taken as cut-off and image binarized to retain only cell bodies by removing smaller objects (Figure 1). The templates were used to quantify Arc levels in NeuN+ or GAD67+ cell bodies. Expression profiles were averaged by normalising cortical thickness via interpolation of images to a height of 6000 pixels (1950 μm, pia to cortical white matter). The mean profile of Arc expression was then compared between delta rhythm and gamma rhythm conditions.

**Figure 1.**
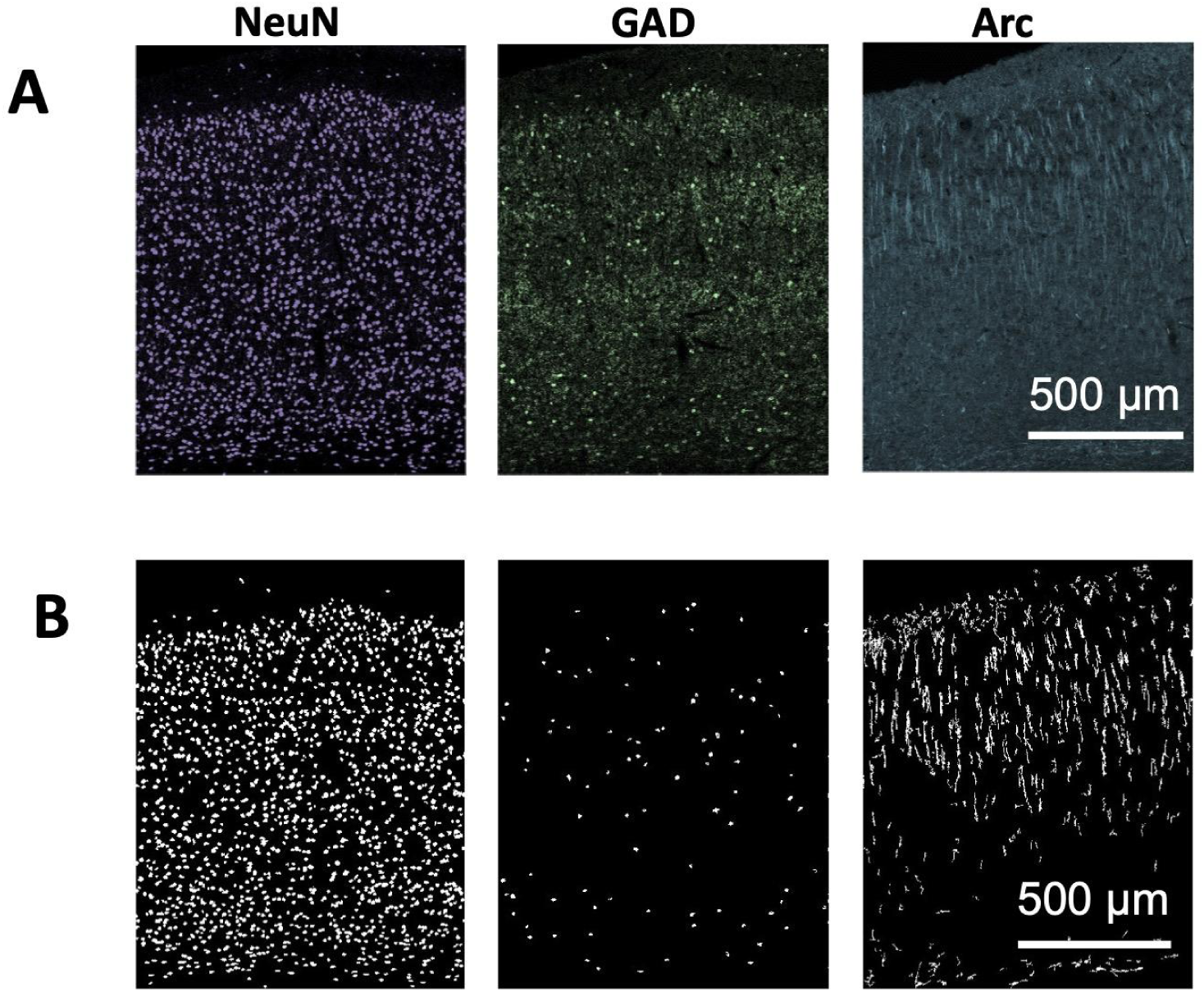
Template formation for image quantification. **A**. Example primary images of NeuN, GAD67 and Arc protein taken form 30 μm sections of secondary somatosensory cortex. **B**. Binary templates generated from each of the fluorescence signals in **A** according to the protocols described in methods. Scale bar 500 μm.

To analyse Arc in dendrites, only the Arc image itself was used. Images were background-subtracted with a rolling kernel of 30 pixels in area. A binary image was then created of the background subtracted image and the threshold used was 1 standard deviation above the mean. To obtain the final dendritic template cell bodies were removed using a ‘roundness’ filter.

## Results

We first compared the overall Arc immunoreactivity in slices that had previously generated sleep-related (delta) or wake-state (gamma) oscillations. Example traces showing neocortical local field potentials of slices treated to evoke oscillations are shown in Figure 2B. The overall expression of Arc was analysed in isolated primary sensory and secondary association cortical regions by measuring the mean fluorescence intensity without consideration for any co-expressed neuronal markers. Figure 2A shows that Arc expression is lower in delta oscillating slices compared to those exhibiting gamma oscillations. In the primary sensory cortex, the mean intensity of Arc staining was lower from 7.85 ± 0.61 in gamma oscillating slices to 5.66 ± 0.56 in delta oscillating slices. In the secondary association cortex the mean intensity of Arc staining was also lower from 7.75 ± 0.67 in gamma oscillating slices to 5.68 ± 0.66 in delta oscillating slices.

**Figure 2.**
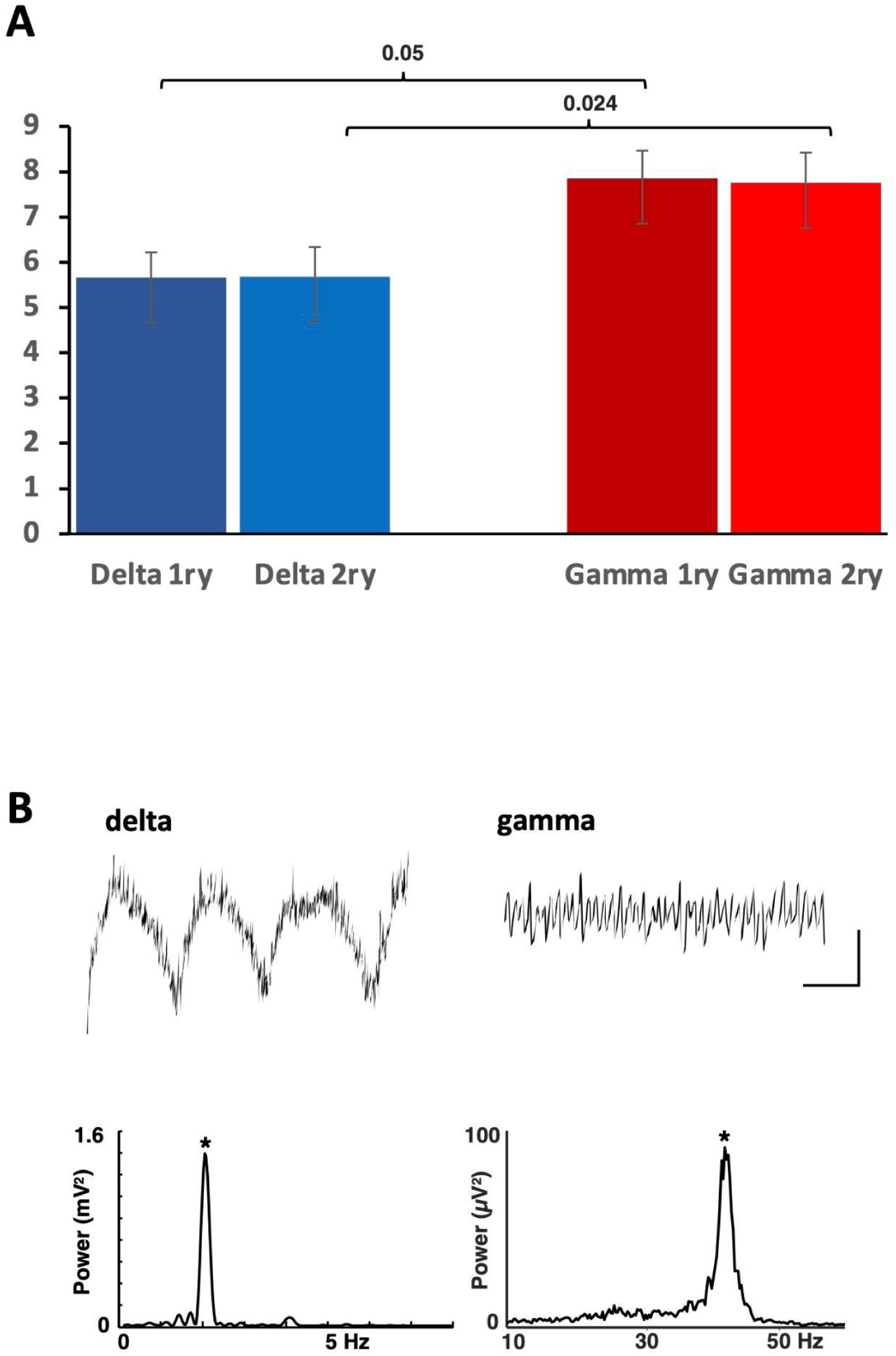
Total Arc immunofluorescence is lower during delta rhythms compared to during gamma rhythms. **A**. Mean total Arc intensity per pixel across the entire primary (1ry) or secondary (2ry) cortical regions studied. (p values are as indicated n=7/6, Student’s t-test). **B.** Example traces showing neocortical local field potentials under the two experimental conditions used: Delta rhythms (left) and gamma rhythms (right). Example spectra are shown for each condition. The power at the modal peak is indicated with asterisks. Scale bars 0.1mV, 200 ms.

Arc is expressed in neuronal nuclei, cell soma and dendritic compartments. To further characterise the distribution of Arc immunoreactivity across the cortex in different cell types and cellular compartments, we next assessed the nuclear and somatic expression of Arc in cells stained either with NeuN, a pan neuronal marker for neuronal nuclei and GAD67, a marker for cortical interneurons. The NeuN and GAD67 binary templates were used to separate nuclear and somatic levels of Arc from dendritic Arc staining and staining in unidentified structures in the neuropil. Arc distribution was assessed in delta and gamma oscillating slices. During both gamma rhythms Arc positive pixels coinciding with NeuN immunoreactivity showed a similar profile across the cortical layers in both the primary and secondary association cortex. The profile was more uniform across the layers in gamma oscillating slices than delta oscillating slices, which showed higher Arc positive pixels in layers 2 and 3. (Figure 3A). There was very little variation in Arc positive pixels in the GAD67+ve cells across the cortical layers and across gamma and delta oscillating slices (Figure 3B).

**Figure 3.**
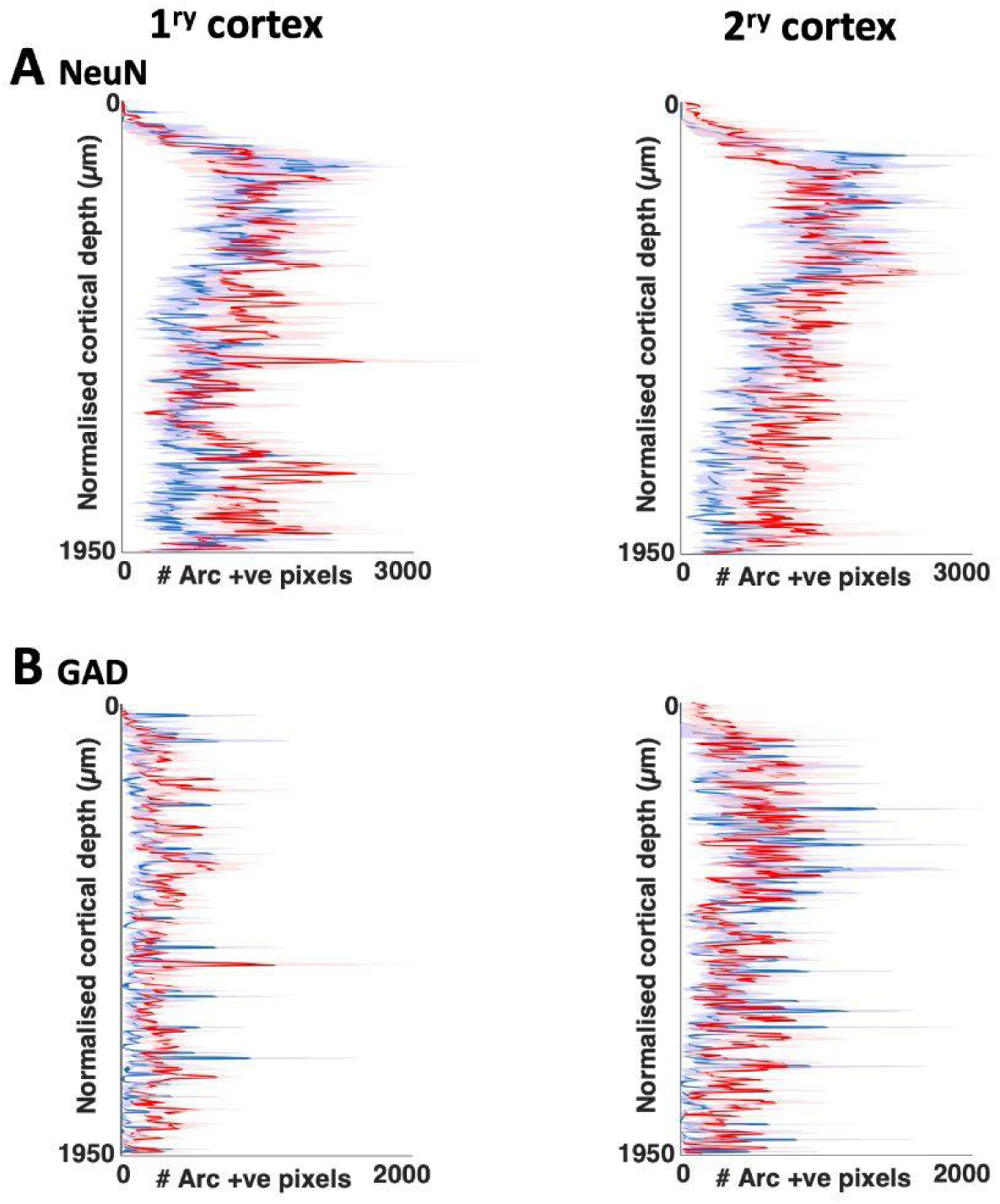
Neuronal somatic Arc protein levels display a uniform pattern across cortical layers after gamma and delta oscillations. **A**. Arc expression in NeuN positive somata was similar in the two rhythms. Graphs show number of Arc immunopositive pixels per tangential scan line coinciding with the NeuN somatic template for primary (left) and secondary (right) neocortical regions in slices generating gamma (red) and delta (blue) rhythms (mean ± s.e.mean, n=7) . **B**. Arc expression in GAD67 positive somata was similar in the two rhythms. Graphs show number of Arc immunopositive pixels per tangential scan line coinciding with the GAD67 somatic template for primary (left) and secondary (right) neocortical regions in slices generating gamma (red) and delta (blue) rhythms.

Across all conditions Arc staining was more prominent in dendrites across the cortical layers (see Figure 4) and an analysis of the profile of dendritic Arc expression across the cortical layers revealed highest Arc positive pixels in layer 2/3 of the primary and secondary association cortex in delta oscillating slices (Figure 5A) . A similar profile was seen in slices showing gamma oscillations so we assessed a correlation between Arc immunoreactivity and rhythm power. Primary sensory cortices and secondary association area images were divided in half tangentially to compare the oscillatory characteristics with Arc expression with cortical layers in superficial and deeper portions of the cortex.

**Figure 4.**
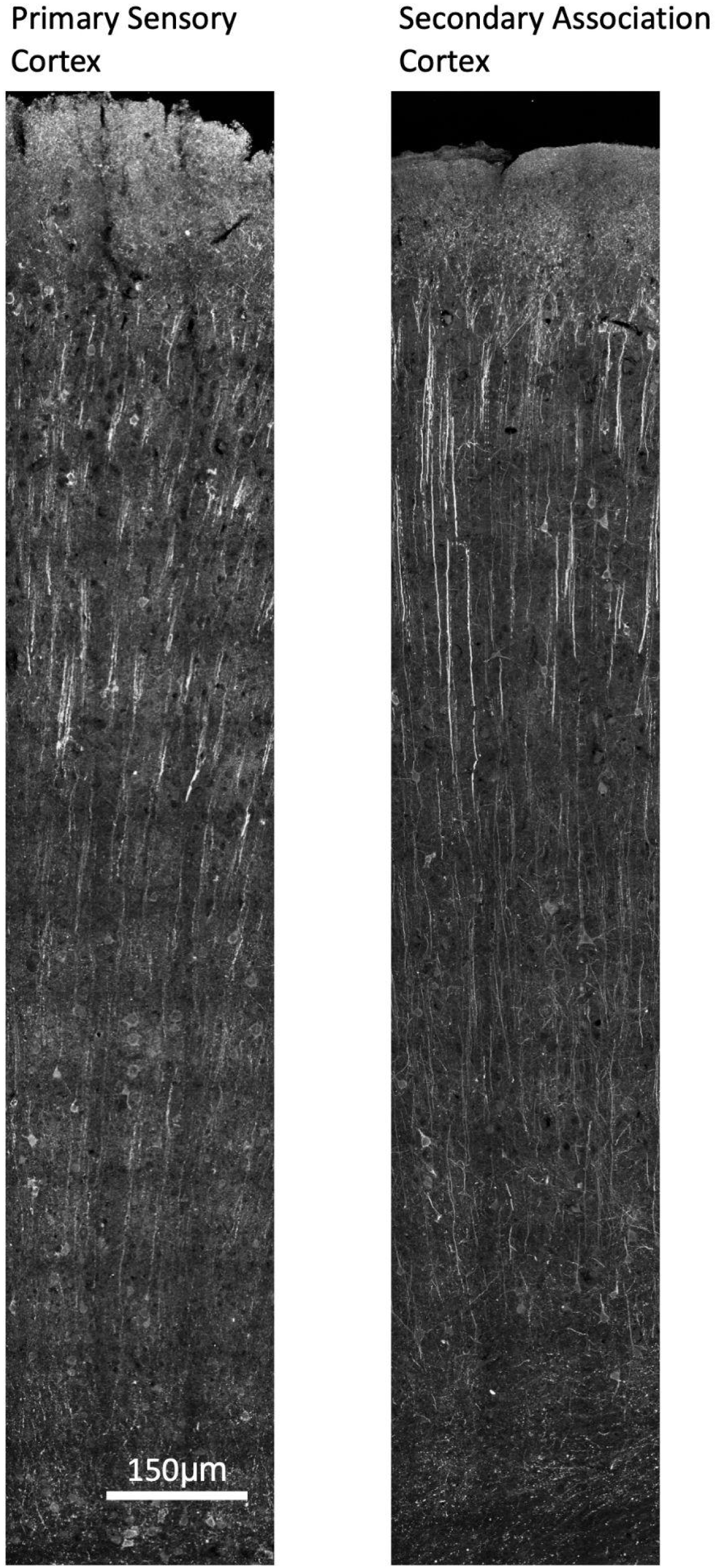
Dendritic Arc staining is more prevalent than somatic staining in neocortical regions. Confocal tile scan images taken at a magnification of 63x from coronal slices of neocortex previously exhibiting delta oscillations in the primary somatosensory cortex or the secondary somatosensory cortex. Scale bar is 150µm.

**Figure 5.**
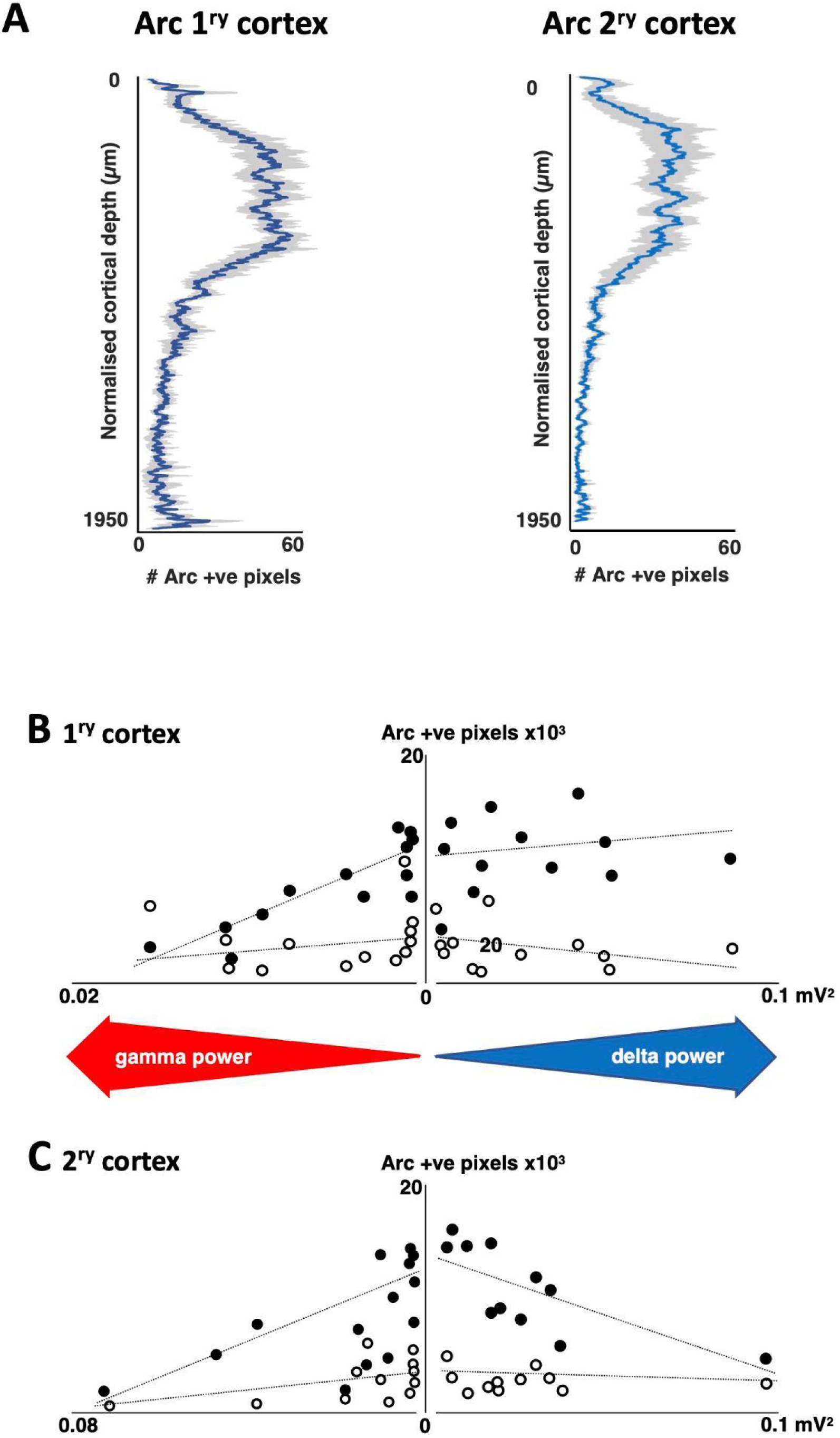
Dendritic Arc protein levels are most prominent in L2/3 dendrites and negatively correlate with gamma power but not delta power. A. Mean number of Arc immunopositive pixels in dendritic compartments are most prominent in L2/3 during delta rhythms. Graphs show number of Arc immunopositive pixels per tangential scan line coinciding with the dendritic template for primary (left) and secondary (right) neocortical regions in slices generating delta rhythms (mean ± s.e.mean, n=7). **B**. Arc immunopositive dendrite expression, plotted against area power, a measure of the rhythm power in primary sensory neocortex. Note the suppression of Arc immunopositive dendrite numbers with increasing gamma power (R^2^ = 0.51) but not increasing delta power. Filled circles indicate superficial layers and open circles indicate deep layers **C**. Arc immunopositive dendrite expression, plotted by rhythm power in secondary, association neocortex. Filled circles indicate superficial layers and open circles indicate deep layers. Note the suppression of Arc immunopositive dendrite numbers in superficial critical layers with both increasing gamma power (linear regression r^2^ = 0.25) and delta power (r^2^ = 0.336).

In the primary sensory cortex (Figure 5B), in delta oscillating slices there was no correlation between the dendritic expression of Arc and delta rhythm power in superficial layers although dendritic expression of Arc was lower in deeper layers than in superficial layers. In contrast, in gamma oscillating slices, superficial layers did display a negative correlation between dendritic Arc and the area power of the oscillation. In the deeper half of the cortex, there were no significant correlations between Arc dendritic staining and the power of gamma oscillations. In the secondary association cortex (Figure 5C), there was a decrease in Arc immunopositive dendrite numbers in superficial critical layers with both increasing gamma power and delta power.

All the data presented thus far has shown that the expression of Arc in dendrites is mostly confined to the superficial layers of the cortex. These dendrites would appear (from visual inspection) to originate in cells whose cell bodies lie in deeper layers and that project their apical dendrites to superficial layers as far as layer 1 (Figure 4).

To identify the specific subtype of the cell responsible for this Arc expression, intracellular electrophysiological recordings were made of single cells, in conjunction with biocytin filling. This allowed for the electrical characterisation of the cells, along with the physical characterisation and co-staining with Arc. This meant that the cell types contributing to the dendritic Arc signal could be revealed. We examined Arc expression layer 5 cell-types that have previously shown to be active during in vitro delta oscillations [21]

Intracellular recordings ruled out layer 5 regular spiking neurons (Supplementary Figure S1) and layer 6 regular spiking neurons (Supplementary Figure S2) as candidates as they did not show co-expression of Arc in their apical dendrite. Intracellular experiments did however successfully show co-staining between biocytin labelled layer 5 intrinsically bursting cells and Arc (Figure 6C).

**Figure 6.**
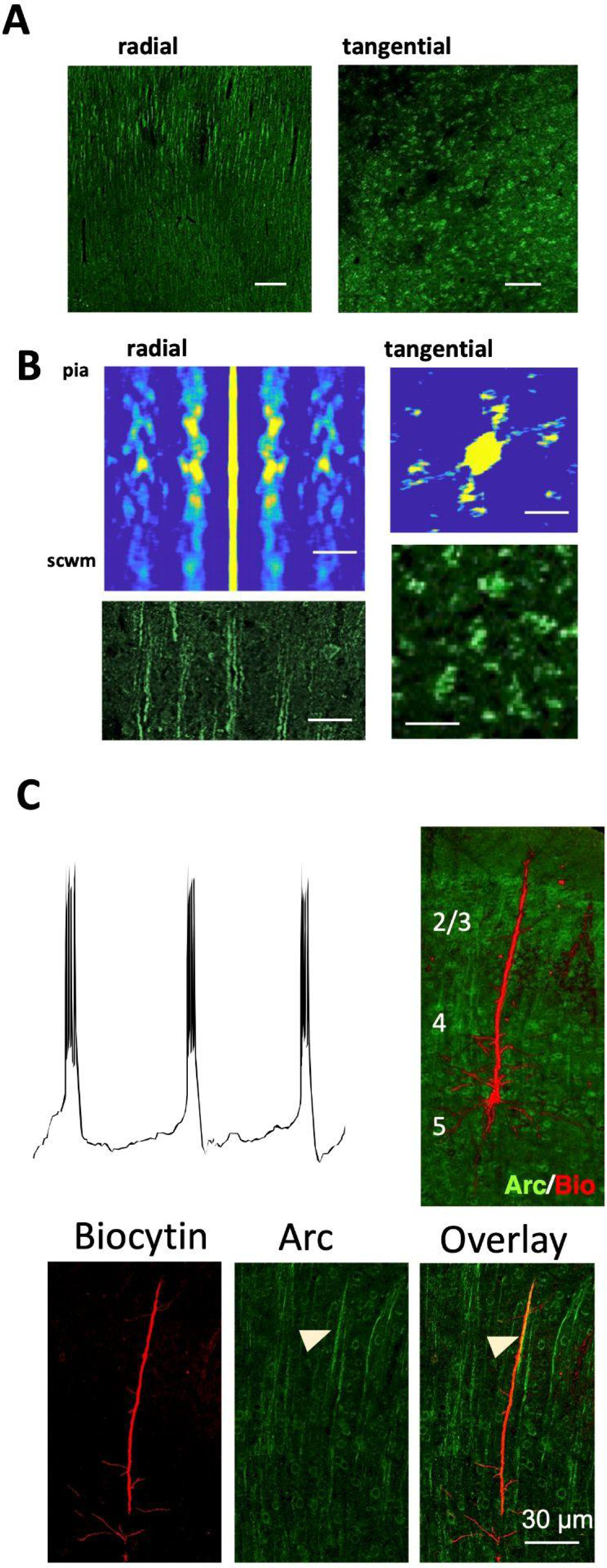
Arc immunopositive dendrites in delta oscillating slices are derived from subcortically-projecting layer 5 pyramidal cells. **A.** Arc immunopositive dendrites were anatomically arranged in clusters, visible predominantly in radial and tangential sections in L2/3. Scale bar 100μm. **B.** Colormaps of mean 1D autocorrelations along scan lines in radial slice images and 2D autocorrelations in tangential slices (cf. Marouka et al., 2017). Images below each autocorrelation show examples of Arc immunofluorescence at the same scale. Scale bars 50 μm. **C**. The only pattern of colocalization of dendrite and Arc was found for primary apical dendrites from biocytin-filled L5 intrinsically bursting neurons. Example trace shows the somatic pattern of burst spiking typical of this neuron subtype during delta rhythms. Right panel illustrates the position of the soma and initial apical dendritic compartments of the neuron used for the example burst behaviour left. Lower panels show higher-magnification images at the level of L2/3 illustrating a more distal apical dendritic compartment from this cell, overlaid with the corresponding Arc signal to show colocalization. Scale bar 30 μm (dendritic images).

The Arc staining was localised to layer 2/3 portion of the apical dendrites of layer 5 intrinsically bursting cells. However, it was also apparent that these Arc immunopositive apical dendrites were not isolated but appeared to be arranged in clusters in radial and tangential sections in layer 2/3 (Figure 6A). To assess this arrangement, a 1-dimensional autocorrelation was performed on each tangential image scan line from pia to subcortical white matter using the original radial sections (Figure 6A). The location of positive pixels of each scan line of the thresholded image was compared with a copy of that line. The copy was then shifted along by one pixel and compared again with itself until there was no overlap. In case of a strong spatial pattern, a high correlation was revealed. The process was repeated for each line of the image and plotted as a colour map. This showed a strong spatially repeating pattern in layers 2/3 with a spatial frequency of ∼50 µm (Figure 6B).

A 2-dimensional autocorrelation analysis was used on tangential slices from the secondary somatosensory cortex to see if this clustering extended beyond the radial plane. We used a similar principle to the autocorrelation mentioned above but involved a rotational comparison. This analysis showed that there was a near-hexagonal structural arrangement of Arc-positive dendritic compartments, again with a spatial frequency of ∼50µm (Figure 6B). The structural arrangement suggests that Arc expression aligns with columnar organisation of layer 5 IB cells.

## Discussion

We have shown that the plasticity-associated protein Arc has a characteristic spatial and cell-type specific distribution during sleep-related delta oscillations in neocortical slices. In delta oscillating slices, Arc is highly expressed in layer 2/3 dendrites of the primary and secondary association cortex and this dendritic localisation is specific to intrinsically bursting cells whose cell bodies are in layer 5. Moreover, Arc immunopositive dendrites are clustered together in a quasi hexagonal arrangement with a spacing of ∼ 50 µm between clusters. We discuss the implications of this complex cytoarchitectural distribution of Arc for the mechanisms of sleep and for the synaptic homeostasis hypothesis regarding the function of sleep.

Our finding that the overall intensity of Arc immunoreactivity in gamma (wake-related) oscillating slices is higher than that in delta (sleep-related) oscillating slices is in line with the previous work [7] that showed a global decrease in Arc in the cerebral cortex during sleep compared to wakefulness. Of note Cirelli and Tonini (2000), found that Arc immunoreactivity increased in dendrites as a function of waking duration - either spontaneous waking or sleep deprivation. Here we found a largely dendritic expression of Arc in both the primary and secondary somatosensory cortex of delta oscillating slices that was highest in layer 2/3 dendrites. If Arc functions in synaptic downscaling via AMPA receptor endocytosis, our data would suggest that synaptic downscaling is more prevalent in particular cortical layers and agrees with the published structural downscaling of layer 2 primary somatosensory synapses [6].

We were able to identify through intracellular recordings, the Arc positive dendrites in delta oscillating slices as apical dendrites of layer 5 intrinsically bursting cells. This laminar and cellular localisation corresponds spatially with calcium spike initiation zones [22], which are portions of the apical dendrite of layer V cells that are highly innervated by inhibitory neurons and as such possess a high density of GABA_B_ receptors. The activation of these receptors has been shown to decrease excitability by the blockage of voltage-dependent calcium channels, which abolishes calcium spikes in these dendrites [23,24]. These metabotropic GABA_B_ receptors are also responsible for the slow prolonged inhibition in layer 2 / 3 of rat somatosensory cortex [25]. Additionally, the GABA_B_ mediated inhibition of layer 5 intrinsically bursting cells is implicated in synaptic depression during delta oscillations and is responsible for the separation of each active phase of the rhythm [21]. The burst firing between periods of inhibition seen during delta rhythms is suggested to be associated with huge calcium influx, which is ideal for the induction of immediate early genes such as Arc. Layer 5 cortical neurons are important for the propagation of delta oscillations [26] and genetic silencing of a subset of layer 5 neurons reduced rebound of slow-wave activity after sleep deprivation [27]. It is notable that Arc knockout mice also show a reduced sleep rebound after sleep deprivation [28].

We found that there was a tangential spatial organisation of clusters of the Arc immunopositive layer 5 IB apical dendrites, at a spatial frequency of ∼50uM. This was conserved when looking at dendritic clusters radially and these structures were also found to have quasi-hexagonal arrangement. Additionally, a within-cluster analysis found this hexagonal-like arrangement to be present between the dendrites of each cluster. Previous work [29] demonstrated a similar feature in the neocortex of mice, with a hexagonal lattice arrangement in the pattern of sub-cortical projection neuron dendrites. The cells that make up these clusters were labelled by the injection of a tracer into the pons and were suggested to represent cortical microcolumns. Not only are layer 5 pyramidal cells known to be anatomically clustered, they also show high c-fos expression [30], and in many areas of the sensory cortex, they fire synchronously with the presentation of stimuli [29]. If the activity of these dendrites is so strongly connected, the high synaptic plasticity between them is unsurprising. The increased change in synaptic plasticity during delta rhythms suggested by the dendritic expression of Arc, most likely related to depression of synapses, occurs in dendrites that form these structures. This suggests that the microcolumns may be the main functional units in the neocortex in which synaptic scaling occurs. It is suggested that whilst in less complex nervous systems single neurons act more independently, the neocortical microcolumn shows that neurons act much synergistically as a single entity in higher animals [31]. Furthermore, there is a high degree of synaptic plasticity between pyramidal neurons in these circuits. Overall, we propose a system whereby synaptic connections are preferentially made and strengthened, as a result of wakeful experiences, in layer 5 pyramidal cells within cortical microcolumns. During the delta oscillations of slow wave sleep, Arc is upregulated, possibly by increased Ca^2+^ entry caused by burst firing of the IB cells. This likely triggers the internalisation of AMPA receptors necessary for synaptic scaling required for the consolidation of memories and the maintenance of synaptic homeostasis.

## Supporting information

Hartnell et al Supplementary figures

## Acknowledgements

We thank members of the Chawla and Whittington Labs for helpful discussion and the Bioscience Technology Facility, University of York for help with confocal imaging. This work was funded by an MRC Case Studentship to IJH.

This paper is dedicated to Professor Miles Whittington who sadly passed away and who was instrumental in the conception and realisation of this study.

Author contributions: IJH, FD and MAW and SC designed research; IJH, AS, SPH and MAW performed research; IJH and MAW analysed data; IJH and SC wrote the paper with editing contributions from all authors.

The authors declare no competing interests.

