## Supplementary material for "Cortical layer-specific and cell type-specific dendritic expression of Arc (Arg 3.1) in an *in vitro* model of slow-wave sleep": Hartnell et al Supplementary figures

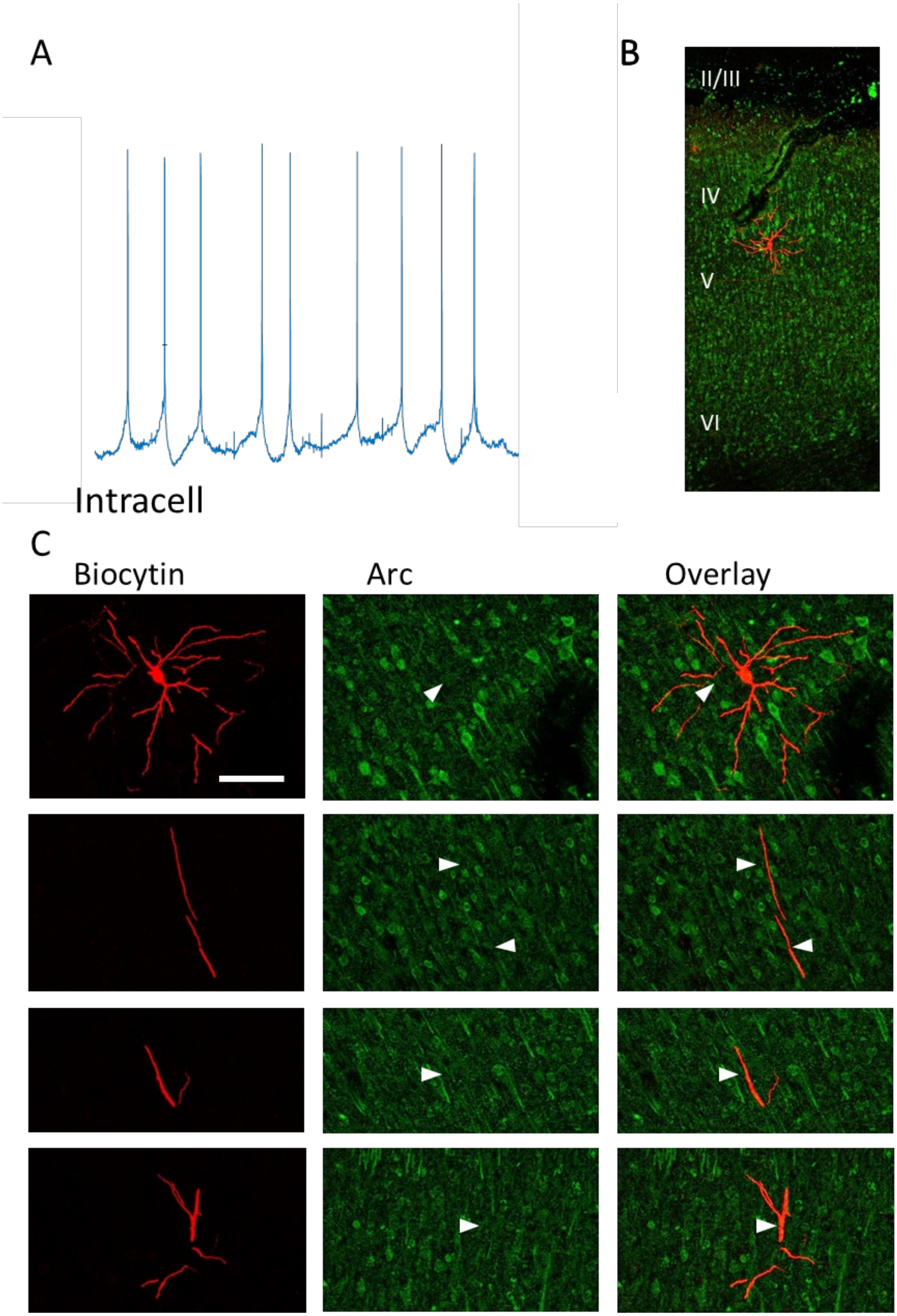


**Figure S1. Arc is not present in the apical dendrites of layer V regular spiking cells in the secondary somatosensory cortex. (A)** 3 second intracellular recording from a layer V regular spiking cell at resting membrane potential. **(B)** Dual stained image of biocytin (red) and Arc (green) from a single sub-slice of the biocytin filled cell, to show the location of the cell body. **(C)** Images of different sections (apical dendrite and cell body) of a layer V regular spiking cell filled with biocytin (red) and co-stained with Arc (green). Scale bar is 100μm. Arrows highlight the location of cell. Cropped images are scale matched.

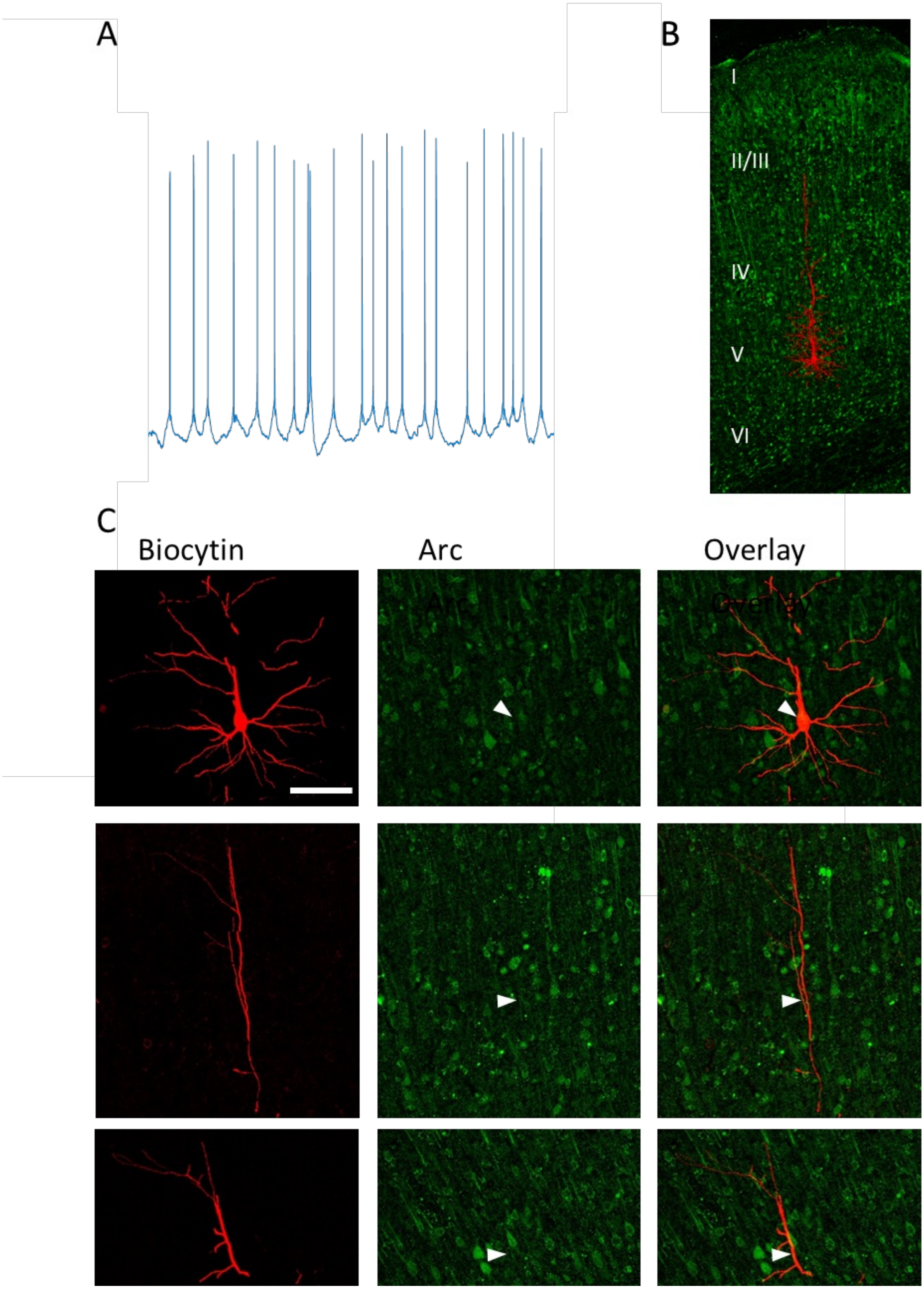


**Figure S2. Arc is not present in the apical dendrites of layer VI regular spiking cells in the secondary somatosensory cortex. (A)** 3 second intracellular recording from a layer VI regular spiking cell at resting membrane potential. **(B)** Dual stained image of biocytin (red) and Arc (green) from a single sub-slice of the biocytin filled cell, to show the location of the cell body. **(C)** Images of different sections (apical dendrite and cell body) of a layer VI regular spiking cell filled with biocytin (red) and co-stained with Arc (green). Scale bar is 100µm. Arrows highlight the location of cell. Cropped images are scale matched.
